# More than noise: Context-dependant luminance contrast discrimination in a coral reef fish (*Rhinecanthus aculeatus*)

**DOI:** 10.1101/2020.06.25.168443

**Authors:** Cedric P. van den Berg, Michelle Hollenkamp, Laurie J. Mitchell, Erin J. Watson, Naomi F. Green, N. Justin Marshall, Karen L. Cheney

## Abstract

Achromatic (luminance) vision is used by animals to perceive motion, pattern, space and texture. Luminance contrast sensitivity thresholds are often poorly characterised for individual species and are applied across a diverse range of perceptual contexts using over-simplified assumptions of an animal’s visual system. Such thresholds are often estimated using the Receptor Noise Limited model (RNL) using quantum catch values and estimated noise levels of photoreceptors. However, the suitability of the RNL model to describe luminance contrast perception remains poorly tested.

Here, we investigated context-dependent luminance discrimination using triggerfish (*Rhinecanthus aculeatus)* presented with large achromatic stimuli (spots) against uniform achromatic backgrounds of varying absolute and relative contrasts. ‘Dark’ and ‘bright’ spots were presented against relatively dark and bright backgrounds. We found significant differences in luminance discrimination thresholds across treatments. When measured using Michelson contrast, thresholds for bright spots on a bright background were significantly higher than for other scenarios, and the lowest threshold was found when dark spots were presented on dark backgrounds. Thresholds expressed in Weber contrast revealed increased contrast sensitivity for stimuli darker than their backgrounds, which is consistent with the literature. The RNL model was unable to estimate threshold scaling across scenarios as predicted by the Weber-Fechner law, highlighting limitations in the current use of the RNL model to quantify luminance contrast perception. Our study confirms that luminance contrast discrimination thresholds are context-dependent and should therefore be interpreted with caution.

## Introduction

The perception of chromatic (colour) and achromatic (luminance) information from the surrounding environment enables animals to perform complex behaviours such as navigation, mate choice, territorial defence, foraging and predator avoidance. Chromatic information is largely used to assess the spectral composition and quality of objects or other organisms (Osorio and Vorobyev, 2005), whereas achromatic information is predominantly used for object grouping, pattern and texture detection, figure-ground segregation, and the perception of motion and depth (Anderson, 2011; Brooks, 2014; Elder and Sachs, 2004; Elder and Velisavljevic, 2010; Gilchrist, 2008; Gilchrist and Radonjic, 2009).

Behavioural experiments to examine colour and luminance discrimination thresholds enable inferences on the perception of visual information by non-human observers (for discussion see Olsson et al., 2018). Thresholds may be influenced by spatiotemporal and spatiochromatic properties of a visual scene, as the perception of colour, pattern, luminance and motion interact when low-level retinal information is processed along pathways in the visual cortex (Monnier and Shevell, 2003; Shapley and Hawken, 2011; Shevell and Kingdom, 2008), or at even earlier stages (Heath et al., 2020; Zhou et al., 2020). For example, the perception of luminance contrast in animals is influenced by a range of factors, including perceived illumination and reflectance (which in turn depend on illumination) in addition to various spatial and temporal properties, such as depth perception, adaptation, stimulus geometry and viewer expectation of the position and shape of a stimulus (Corney and Lotto, 2007; Craik, 1938; Gilchrist and Radonjic, 2009; Heinemann and Chase, 1995; Kingdom, 2011; Lind et al., 2012; Pelli and Bex, 2013). The impact of post-photoreceptor, and particularly post-retinal neuronal processing, on luminance perception is often illustrated by visual displays targeting these effects, such as simultaneous contrast illusions (Fig. 1). To investigate the design, function and evolution of animal visual signals, the context sensitivity of visual threshold measurement is important to define.

**Figure 1:**
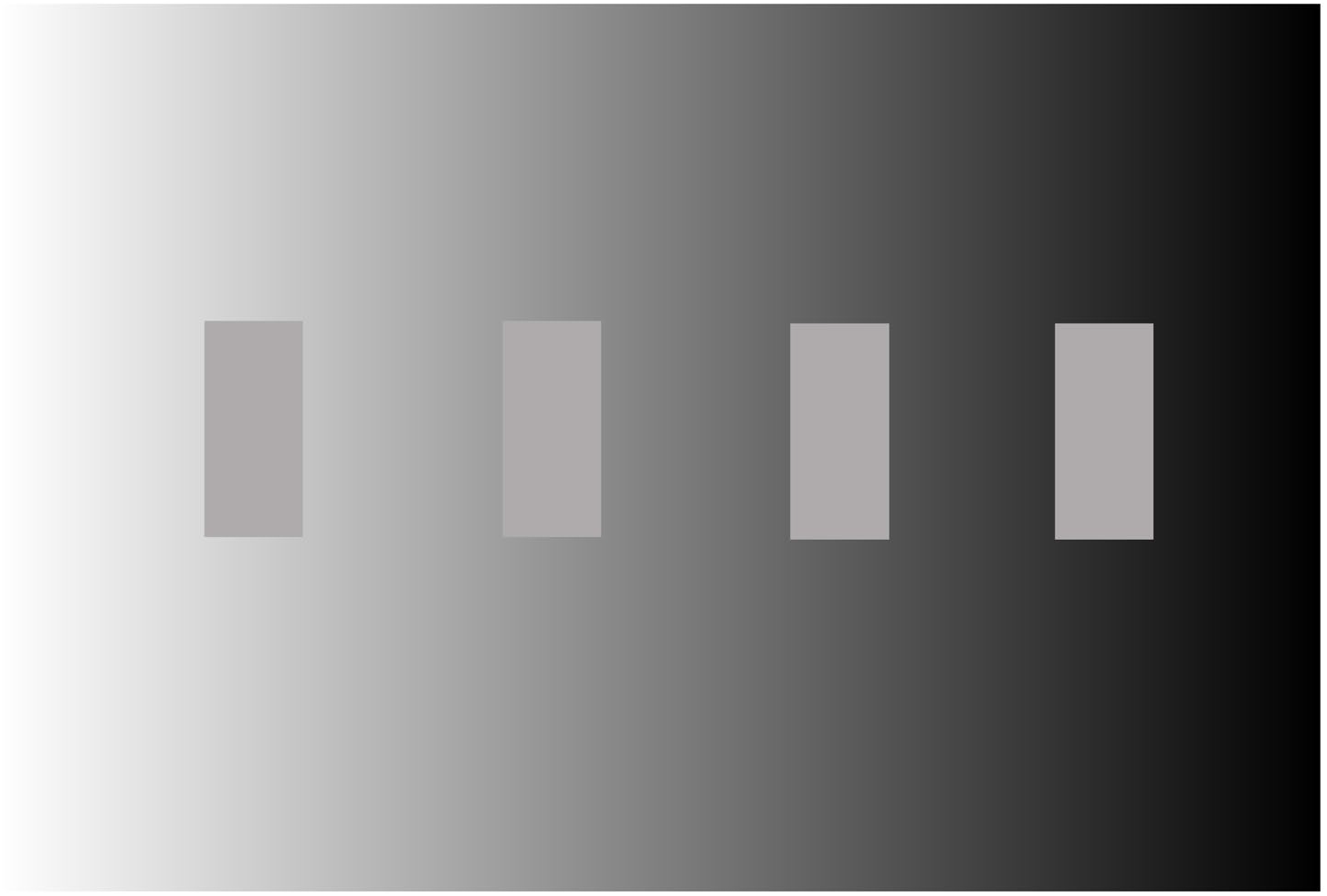
The simultaneous contrast effect: despite having identical luminance, the left most internal square appears darker than the right one as a result of the background contrast against which they each square is viewed.

Luminance contrast of objects against their visual background or between objects can be measured in a number of different ways, including Michelson contrast (MC), Weber contrast (WC) and Root Mean Square (RMS) (Bex and Makous, 2002; Moulden et al., 1990; Vorobyev and Osorio, 1998). MC is commonly used to describe the contrast between two comparably sized objects or sine gratings (Bex and Makous, 2002; Pelli and Bex, 2013). The WC, particularly popular in psychophysics, is designed to describe the contrast of an object against a dominating background, while accounting for the Weber-Fechner law that states that psychometric thresholds scale with stimulus intensity at a constant ratio: the Weber fraction (Dzhafarov and Colonius, 1999; Norwich, 1987; Treisman, 1964). Luminance discrimination thresholds in animals have been obtained from behavioural experiments and measured in MC, and most commonly in WC (e.g. Lind et al., 2013; Scholtyssek et al., 2008). For example, human luminance discrimination thresholds are between 0.11 and 0.14 WC (Cornsweet and Pinsker, 1965), which is similar to seals (0.11-0.14 WC) (Scholtyssek and Dehnhardt, 2013; Scholtyssek et al., 2008). Other animals have poorer luminance discrimination thresholds, including birds (0.18-0.22 WC) (Lind et al., 2013), dogs (0.22-0.27 WC) (Pretterer et al., 2004), manatees 0.35 WC (Griebel and Schmid, 1997) and horses (0.42-0.45 WC) (Geisbauer et al., 2004).

Behavioural experiments measuring discrimination thresholds are often time-consuming and unfeasible, especially when studying non-model organisms. Furthermore, focal species may not be suitable for behavioural testing due to ethical, legal or logistical restrictions. Therefore, in studies on visual ecology, the ‘Receptor Noise Limited’ (RNL) model (Vorobyev and Osorio, 1998) has been adopted as a means of estimating whether both colour and luminance contrast within and between animal colour patterns, or between animals and their backgrounds, are perceivable to a species The model was initially designed for colour contrast modelling; however, the achromatic interpretation of the RNL model (Siddiqi, 2004).has been used in a large number of studies to quantify the perception of luminance contrast by non-human observers (e.g. Cheney and Marshall, 2009; Marshall et al., 2016; Spottiswoode and Stevens, 2010; Stoddard and Stevens, 2010; Troscianko and Stevens, 2015). In contrast to using WC or MC, the RNL model allows the prediction of contrast discriminability without the need of behavioural experimentation, but instead can even be applied using conservatively chosen estimates of vision parameters (Olsson et al., 2018).

The RNL model assumes that signal discrimination under ‘ideal viewing conditions’ is limited by noise originating in the receptors and subsequent opponent processing (Vorobyev and Osorio, 1998; Vorobyev et al., 2001). It was designed to estimate when a signal receiver could discriminate between two colours that were spectrally similar, adjacent, of fixed size and luminance. The point at which the contrast between two stimuli surpasses a behaviourally determined threshold (e.g. 75% correct choice in a pairwise choice paradigm) is then expressed as a ‘Just Noticeable Difference’ (JND) corresponding to a Euclidian distance (ΔS) in an *n*-dimensional space, where *n* is the number of colour or luminance processing channels (Hempel de Ibarra et al., 2001). The model predicts a JND is equal to 1 ΔS if all model assumptions (ideal viewing conditions) are met (Vorobyev and Osorio, 1998; Vorobyev et al., 2001).

However, in many animals, the neuronal pathways leading to the perception of luminance contrast vary significantly from those involved in the perception of colour contrast. In humans for example, the magnocellular and parvocellular pathways segregate colour and luminance tasks (Zeki, 1993) which can interact (to varying degrees) during subsequent neuronal processing (e.g. Bruce et al., 2010; Gegenfurtner and Kiper, 1992; Shapley and Hawken, 2011; Simmons and Kingdom, 2002; Webster and Wilson, 2000). The pronounced context-dependent sensitivity of luminance contrast perception is partly due to the fact that achromatic vision in vertebrates lacks a process as efficient as colour constancy (Kelber et al., 2003; Land, 1986; Osorio and Vorobyev, 2008; Wallach, 1948) which enables the perceived color of objects to remain relatively constant under varying illumination conditions (but see Lotto and Purves, 2000; Simpson et al., 2016). However, despite assuming receptor noise levels to be the limiting factor shaping both chromatic and achromatic contrast perception, behavioural validations of perceptual distances calculated using the RNL model are required in various visual contexts (as suggested by Olsson, Lind, & Kelber, 2018 but see Skorupski & Chittka, 2011; Vasas, Brebner, & Chittka, 2018). Olsson et al. (2018) have further suggested a conservative threshold of up to 1 JND = 3 ΔS for colour discrimination, as both parameter choice and behavioural threshold validation are often difficult. The use of such conservative chromatic discrimination thresholds in perceptually complex contexts has recently been supported by empirical work (Escobar-Camacho et al., 2019; Sibeaux et al., 2019). However, no empirical evidence exists for choosing conservative luminance (achromatic) contrast thresholds using the RNL model.

In this study, we performed behavioural experiments with triggerfish, *Rhinecanthus aculeatus*, to determine luminance discrimination thresholds in a foraging task using large stimuli under well-illuminated (photopic) conditions. We refer to the task of discriminating a stimulus from its background as a detection task, as this reflects a common use of the achromatic RNL model in visual ecology, most prominently when quantifying the efficiency of animal camouflage (e.g. Troscianko *et al.*, 2016). The ability to detect the presence of a potential prey item is the pre-requisite for more complex cognitive processes and decision making by a predator (Endler, 1991) and as such more likely to reflect low-level retinal and post-retinal properties of visual contrast processing, such as the ones the RNL model has been developed to reflect.

Fish were trained to first locate a target spot that was randomly placed on an achromatic background from which the spot differed in terms of luminance, and then peck it to receive a food reward. Luminance discrimination thresholds were measured for both increasing and decreasing luminance, on both a relatively bright and a dark background. We report thresholds in terms of Michelson and Weber contrast, but then translate these thresholds into achromatic ΔS using the log transformed RNL model, as per Siddiqi *et al.* (2004). To our knowledge, this is the first time that achromatic discrimination thresholds have been quantified in a marine vertebrate, using a ‘detection’ task (as opposed to a pairwise choice paradigm as in Siebeck et al., (2014)) as well as doing so using animals which have been trained to detect both randomly placed brighter and darker stimuli simultaneously.

## Materials and methods

### Study species

We used triggerfish *Rhinecanthus aculeatus* (n = 15), which ranged in size from 6 to 16 cm (standard length, SL). This species inhabits shallow tropical reefs and temperate habitats throughout the Indo-Pacific and feeds on algae, detritus and invertebrates (Randall et al., 1997). They are relatively easy to train for behavioural studies (e.g. Green *et al.*, 2018), and their visual system has been well-studied (Champ et al., 2014; Champ et al., 2016; Cheney et al., 2013; Pignatelli et al., 2010). They have trichromatic vision based on one single cone, containing short-wavelength sensitive visual pigment (sw photoreceptor λ_max_ = 413 nm); and a double cone, which houses the medium-wavelength sensitive pigment (mw photoreceptor λ_max_ = 480 nm) and long-wavelength sensitive pigment (lw photoreceptor λmax = 528 nm) (Cheney et al., 2013). The double cone members are used independently in colour vision (Pignatelli et al., 2010), but are also thought to be used in luminance vision (Marshall et al., 2003; Siebeck et al., 2014), as per other animals such as birds and lizards (Lythgoe, 1979).

However, it is not clear if both members of the double cone are used for luminance perception via electrophysiological coupling (Marchiafava, 1985; Siebeck et al., 2014). We have based our experiment on the assumption of both members contributing as per previous studies modelling luminance perception in *R. aculeatus* (Mitchell et al., 2017; Newport et al., 2017). These studies have used the added input of both double cone members (mw + lw), whereas our study uses the averaged output of both members (mw + lw / 2) as suggested by Pignatelli & Marshall, (2010) and Pignatelli et al., (2010). Additionally, Cheney et al., (2013) have used the lw receptor response rather than both double cone members for luminance contrast modelling in *R. aculeatus*, based on discussions in Marshall et al., (2003). However, MC/WC/ΔS contrast values are identical for f_t/b_ = mw + lw and f_t/b_ = mw + lw / 2 (eq. 2). Using the lw member of the double cone only (as opposed to both members) causes less than 1% difference (well below measurement error) in receptor stimulation due to the lack of chromaticity of the stimuli and the strong overlap of spectral sensitivities of both double cone members (Cheney et al., 2013).

Fish were obtained from an aquarium supplier (Cairns Marine Pty Ltd, Cairns), shipped to The University of Queensland, Brisbane and housed in individual tanks of 120L (W: 40cm; L: 80cm, H: 40cm). They were acclimatised for at least one week before training commenced. Experiments were conducted in September-November 2017. All experimental procedures for this study were approved by the University of Queensland Animal Ethics Committee (SBS/111/14/ARC).

### Stimulus creation and calibration

We used a custom programme in Matlab (MathWorks, 2000) to create the stimuli (available on GitHub). This programme allowed us to specify the RGB values of the background and target spot, and randomly allocate the target spot (1.6cm diam) to a position on the background. The size of spot was chosen to be well within the spatial acuity of *R. aculeatus* (Champ et al., 2014) and could be easily resolved by the fish from anywhere in their aquaria. Stimuli, distractors and backgrounds were printed on TrendWhite ISO 80 A4 recycled paper using a HP Laserjet Pro 400 color M451dn printer. Stimuli were then laminated using matte laminating pouches. Throughout the experiment, any stimuli with detectable scratches or damage were replaced immediately.

To ensure all stimuli were achromatic, reflectance measurements were plotted in colour space as per Cheney et al. (2019). Target and background colours were < 1 ΔS from the achromatic locus in the RNL colour space as per equations 1-4 in Hempel de Ibarra et al. (2001). Photoreceptor stimulation was calculated using spectral sensitivities of triggerfish from Cheney et al. (2013). Measures of photoreceptor noise are not available in this species, therefore we assumed a cone ratio of 1:2:2 (SW:MW:LW) with a standard deviation of noise in a single cone of 0.05 as per (Champ et al., 2016; Cheney et al., 2019). The cone abundance was normalised relative to the LW cone, which resulted in channel noise levels (univariant Weber fractions) of 0.07:0.05:0.05 (SW:MW:LW).

We quantified luminance contrast using calibrated digital photography (Stevens et al., 2007) using an Olympus E-PL5 Penlight camera fitted with a 60mm macro lens to take pictures of each stimulus combination (Suppl. material). Two EcoLight KR96 30W white LED lights (Eco-lamps Inc. – Hong Kong) were used to provide even illumination between 400-700nm wavelength (Suppl. Material). Pictures were analysed using the ‘Multispectral Image Calibration and Analysis’ (MICA) Toolbox (Troscianko and Stevens, 2015) to calculate cone capture quanta of the double cone. The double cone stimulation was calculated as the average stimulation of the medium-wavelength (MW) and long-wavelength (LW) cone, as per Pignatelli *et al.* (2010). We used a spatial acuity estimation of 2.75 cycles per degree (Champ et al., 2014) at 15cm viewing distance using AcuityView (Caves and Johnsen, 2018) implemented in MICA’s QCPA package (van den Berg et al., 2020).

Stimulus contrast was measured as Michelson contrast using the MICA derived cone catch values of the double cones. The stimuli contrasts were evenly spaced around an area of interest in which the threshold was expected to lie, according to pilot trials. Weber contrast of the thresholds was calculated as Δ*I_t_*/*I_s_*; where Δ*I_t_* is the stimulus contrast at threshold and *I_s_* is the intensity of the distractor or background respectively as per Lind *et al.*, (2013). Achromatic ΔS values were calculated according to equation 7 in Siddiqi *et. al* (2004) (Eq. 1).

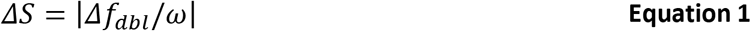

Where Δ*f*_*dbl*_ describes the contrast in von Kries corrected double cone stimulation between the stimulus (*f*_*t*_) and its background (*f*_*b*_), calculated as per equation 4 in Siddiqi *et. al* (2004) (Eq. 2) in relation to the weber fraction (*w*) of the double cone channel. When using the natural logarithm of the quantum catches *w* = *e*_*i*_

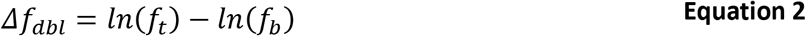

A total of 6 stimuli were created for each scenario (Fig. 2, Table 1).

**Figure 2:**
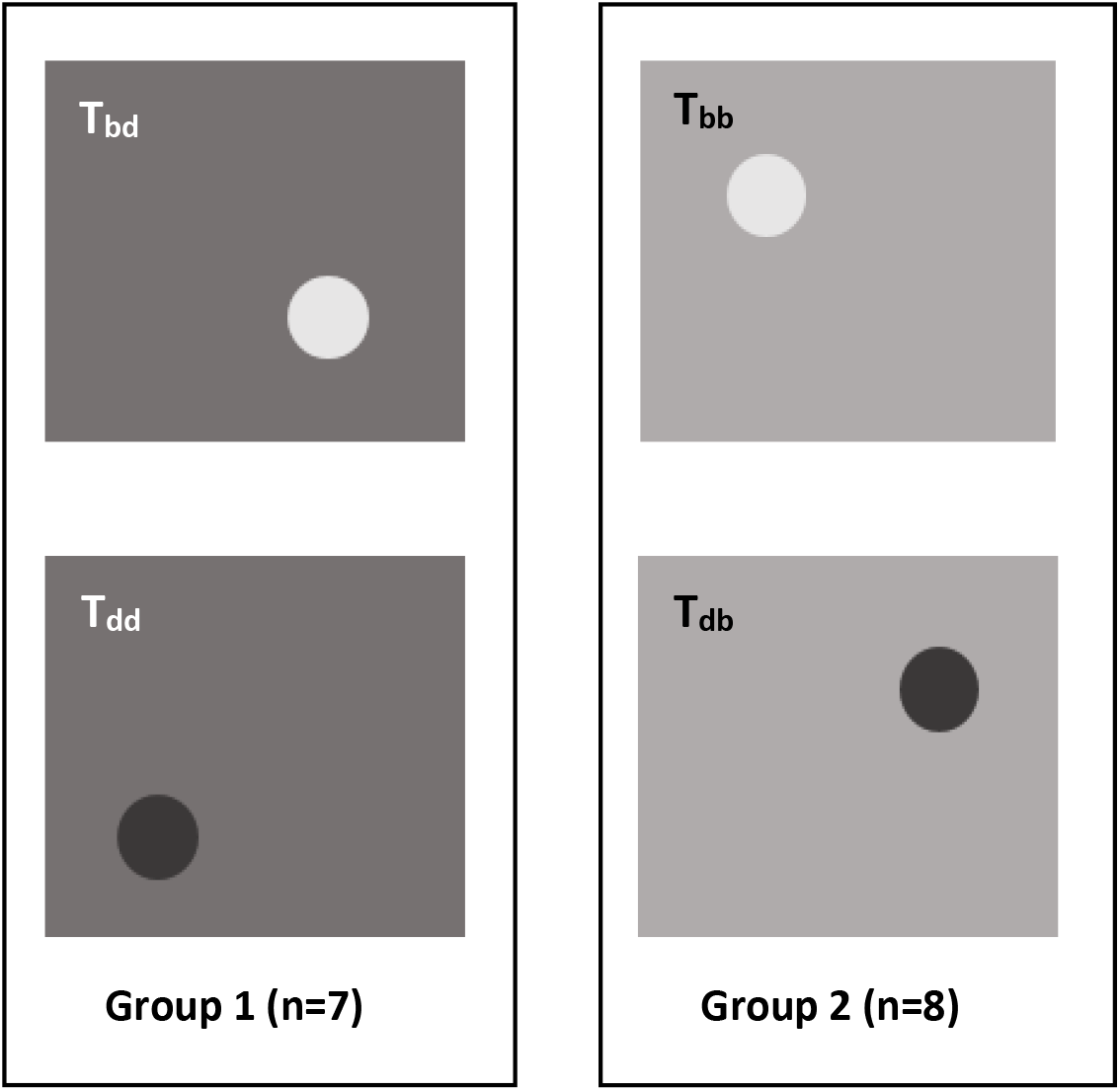
Schematic representation of detection scenarios. Figure proportions are not to scale. Group 1 (dark background) and group 2 (bright background) are shown with darker and brighter target spots with the maximum contrast used in the experiment. The top left of each scenario shows the corresponding abbreviation. T_bd_ = bright spot on dark background, T_dd_ = dark spot on dark background, T_bb_ = bright spot on bright background, T_db_ = dark spot on bright background. Background were A4 size and the spots 1.6cm in diameter, randomly placed of each trial.

**Table 1:** Summary of all stimulus contrasts across both groups in ΔS and Michelson contrast. Number of trials per fish are indicated in brackets below each stimulus contrast.

| Group 1 (dark background, n = 7) |  | Group 2 (bright background, n = 8) |  |
| --- | --- | --- | --- |
| [ΔS] / [Michelson Contrast] |  | [ΔS] / [Michelson Contrast] |  |
| (median trials per fish ± sd) |  | (median trials per fish ± sd) |  |
| Bright Spot (T<sub>bd</sub>) | Dark Spot (T<sub>dd</sub>) | Bright Spot (T<sub>bb</sub>) | Dark Spot (T<sub>db</sub>) |
| 15.34 / 0.37* | 9.26 / 0.23* | 17.87 / 0.42* | 15.51 / 0.37* |
| (6.0 ± 0.5) | (7.0 ± 1.1) | (8.5 ± 1.5) | (9.0 ± 1.8) |
| 5.98 / 0.15 | 6.55 / 0.16 | 8.84 / 0.22 | 7.99 / 0.20 |
| (8.0 ± 1.3) | (6.0 ± 0.4) | (8.5 ± 1.8) | (8.0 ± 1.3) |
| 4.82 / 0.12 | 5.04 / 0.13 | 5.19 / 0.13 | 5.92 / 0.15 |
| (6.0 ± 0.5) | (8.0 ± 1.7) | (7.5 ± 0.9) | (8.5 ± 1.5) |
| 3.94 / 0.10 | 3.03 / 0.08 | 3.98 / 0.10 | 4.65 / 0.12 |
| (8.0 ± 1.5) | (9.0 ± 1.5) | (9.0 ± 1.7) | (8.5 ± 1.3) |
| 2.34 / 0.06 | 1.24 / 0.03 | 1.82 / 0.05 | 2.46 / 0.06 |
| (8.0 ± 1.2) | (9.0 ± 1.6) | (8.0 ± 1.3) | (6.0 ± 1.6) |
| 0.58 / 0.01 | 0.89 / 0.02 | 0.84 / 0.02 | 1.58 / 0.04 |
| (7.0 ± 1.1) | (9.0 ± 1.6) | (6.5 ± 1.0) | (7.0 ± 1.4) |

### Experimental setup

Aquaria were divided in two halves by a removable grey, opaque PVC partition. This enabled the fish to be separated from the testing arena while the stimuli were set up. Stimuli were displayed on vertical, grey, PVC boards and placed against one end of the aquaria. Tanks were illuminated using the same white LED lights (EcoLight KR96 30W) used for stimulus calibration. To ensure equal light levels in all tanks, sidewelling absolute irradiance was measured using a calibrated OceanOptics USB2000 spectrophotometer, a 180° cosine corrector and a 400nm optic fibre cable fixed horizontally in the tank (Suppl. Material).

### Animal training

Fish were trained to peck at the target dot using a classic conditioning approach. First, fish were trained to pick a small piece of squid off a black or white (randomly chosen) spot (1.6 cm diam) on the grey background corresponding to the treatment group (‘bright’ or ‘dark’, Table 1). We trained the fish to detect target spots on both brighter and darker backgrounds to reduce hypersensitivity through anticipation by applying the principle of ‘constant stimuli’ thresholds (Colman, 2008; Laming and Laming, 1992; Pelli and Bex, 2013). We will be referring to stimuli with greater luminance than their background as bright or brighter to facilitate reading. However, the perception of luminance is complex, and the term brightness means specifically the perception of surface luminance is often used wrongly and/or in confusion with lightness which refers to the perception of surface reflectance (Kingdom, 2011). Training fish to react to stimuli being either brighter or darker intended to produce thresholds more closely related to a natural context, as prey items in the natural environments can be both brighter and darker than their natural background. Second, once fish consistently removed the food reward from the black and white target spots, a second food reward was presented from above using forceps. Once fish were confident with this, the final stage of training was a food reward given from above once they had tapped at the target stimulus (without food). Training consisted of up to two sessions per day, with six to ten trials per session. Fish moved to the testing phase when fish were successful in performing the task in > 80% trials over at least 6 consecutive sessions. A trial was considered unsuccessful if the fish took longer than 90 seconds to make a choice or if it pecked at the background more than twice. Testing was suspended for the day if the fish showed multiple timeouts for obviously easy contrasts, assuming the fish was not motivated to perform the task. However, this occurred rarely (<1% of trials) with smaller fish being more susceptible to having been fed enough to lose appetite.

### Animal testing

We randomly allocated fish into two groups: Group 1 (n = 7) had to find and peck at target spots that were brighter (T_bd_) or darker (T_dd_) than a relatively dark background; Group 2 (n = 8) had to find and peck target spots that were brighter (T_bb_) or darker (T_db_) than a relatively bright background (Fig. 2, Table 1). As with the training of the animals, the target spots were presented in a random position against an A4 sized achromatic background in two sessions per day consisting of 6-10 trials per session depending on the appetite of the fish. The trials for each session were chosen pseudo-randomly from all possible contrasts, thus fish were presented with both darker and brighter spots compared to their background in each session. Each stimulus was presented a minimum of 6 times (Table 1). We ensured that both easier and harder contrast stimuli were presented in each session to maintain fish motivation. Motivation was considered low when the animal did not engage in the trial immediately, and if this occurred, trials were ceased for that fish until the next session. However, this rarely occurred and was further minimised by carefully avoiding overfeeding the animals. As per training, trial was considered unsuccessful if the fish took longer than 90 seconds to make a choice or if it pecked at the background more than twice. Wrong pecks were recorded and time to detection was recorded as the time between the moment the fish moved past the divider and the successful peck at the target spot.

### Statistical analysis

Psychometric curves were fitted to the data with % correct choice per stimulus as the response variable and stimulus contrast measured in Michelson contrast as the independent variable, using the R package *quickpsy* (Linares and Lopez-Moliner, 2015; R Core Team, 2015). The best model fit (cumulative normal or logistic) was determined using the lowest AIC as per Yssaad-Fesselier & Knoblauch (2006) and Linares & Lopez-Moliner (2015) and is expressed both individually for each scenario as well as the sum across all scenarios. We interpolated the 50% correct choice thresholds with a 95% confidence interval from these curves. Thresholds between the fitted curves for each scenario were compared as per Jörges *et al.* (2018) using the Bootstrap (Boos, 2003) implemented in *quickpsy* (100 permutations). The Bonferroni method (Bland and Altman, 1995) was used to adjust the significance level of the confidence intervals to 1-0.05/n, with n corresponding to the number of comparisons.

## Results

A total of 1365 trials were conducted across all animals and treatments (Table 1). The total success rate was 68.5% across all 24 stimuli with a median (± sd) time to detection of 3.1 ± 12.6 s with the fastest success at 0.3 seconds and the slowest at 89.9 s. The median time for successful detection was similar across all scenarios (± sd): T_dd_ = 2.9 ± 12.9 s, T_bd_ = 2.8±10.8 seconds, T_db_ = 3.1 ± 13.5 s, T_bb_ = 3.22 ± 12.58 s.

Detection thresholds (50% correct choice) for all scenarios are presented in Figure 3 and Table 2. The sum of AIC across all four detection scenarios (fit = cumulative normal) was 162.4 (T_dd_ = 24.2, T_bd_ = 50.8, T_bb_ = 50.1, T_db_ = 37.3). In group 1 (dark background), the detection thresholds for the bright and dark spot were not significantly different from each other, with the threshold for detecting a spot brighter than a dark background being slightly higher than a spot darker than a dark background (T_bd_ - T_dd_ = 0.007 MC, CI_diff_ [0.002 / 0.017]). However, the detection thresholds in group 2 (bright background) were significantly different from each other, with the threshold for detecting a dark spot against a bright background being significantly lower than the threshold for detecting a bright spot against a bright background (T_db_ - T_bb_ = −0.028MC, CI_diff_ [0.014 / 0.041]).

**Figure 3:**
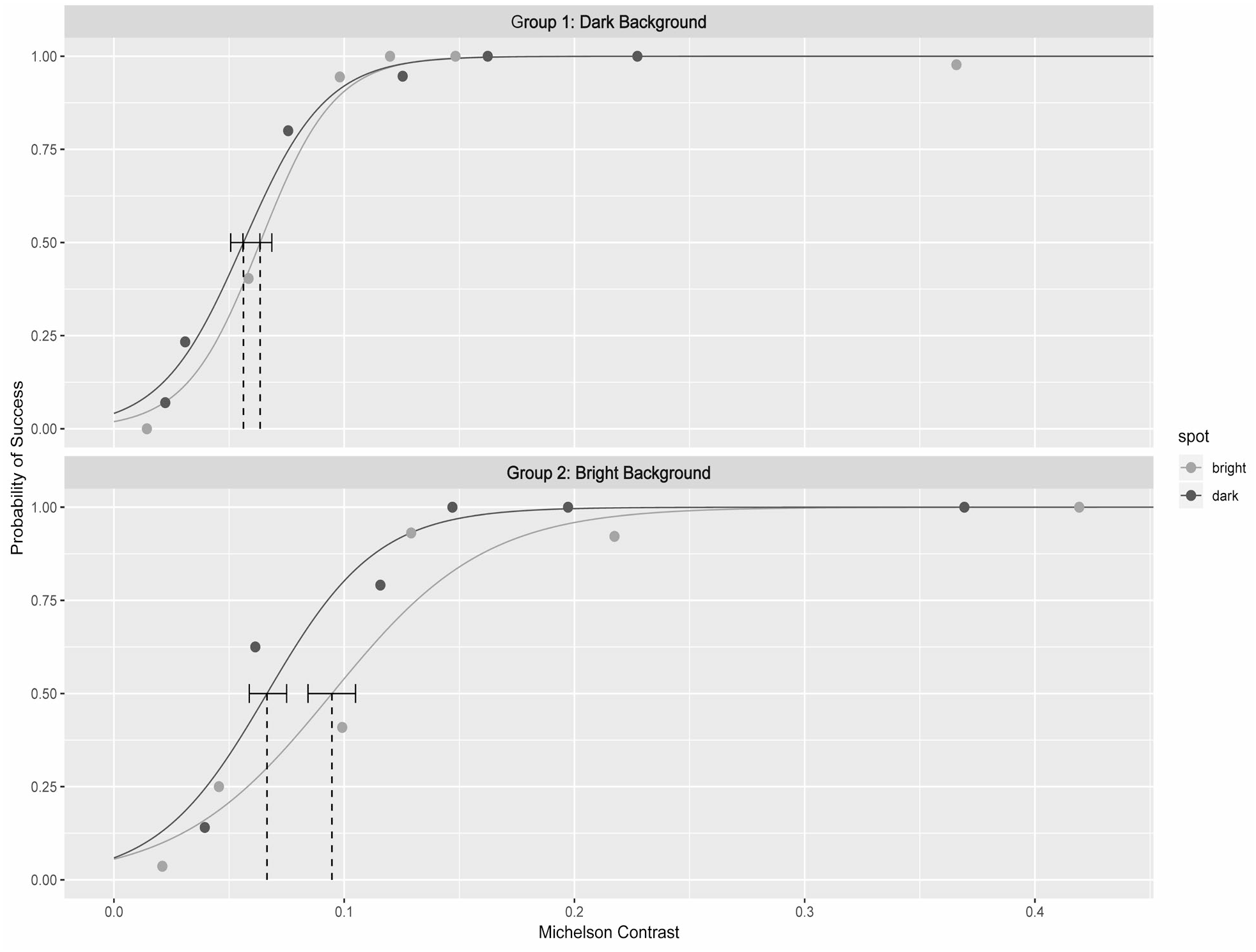
50% probability of a fish successfully pecking a target spot. Estimated using a logistic regression fitted to the detection data. Thresholds for each scenario in Michelson contrast, error bars represent the 95% confidence intervals.

**Table 2:**
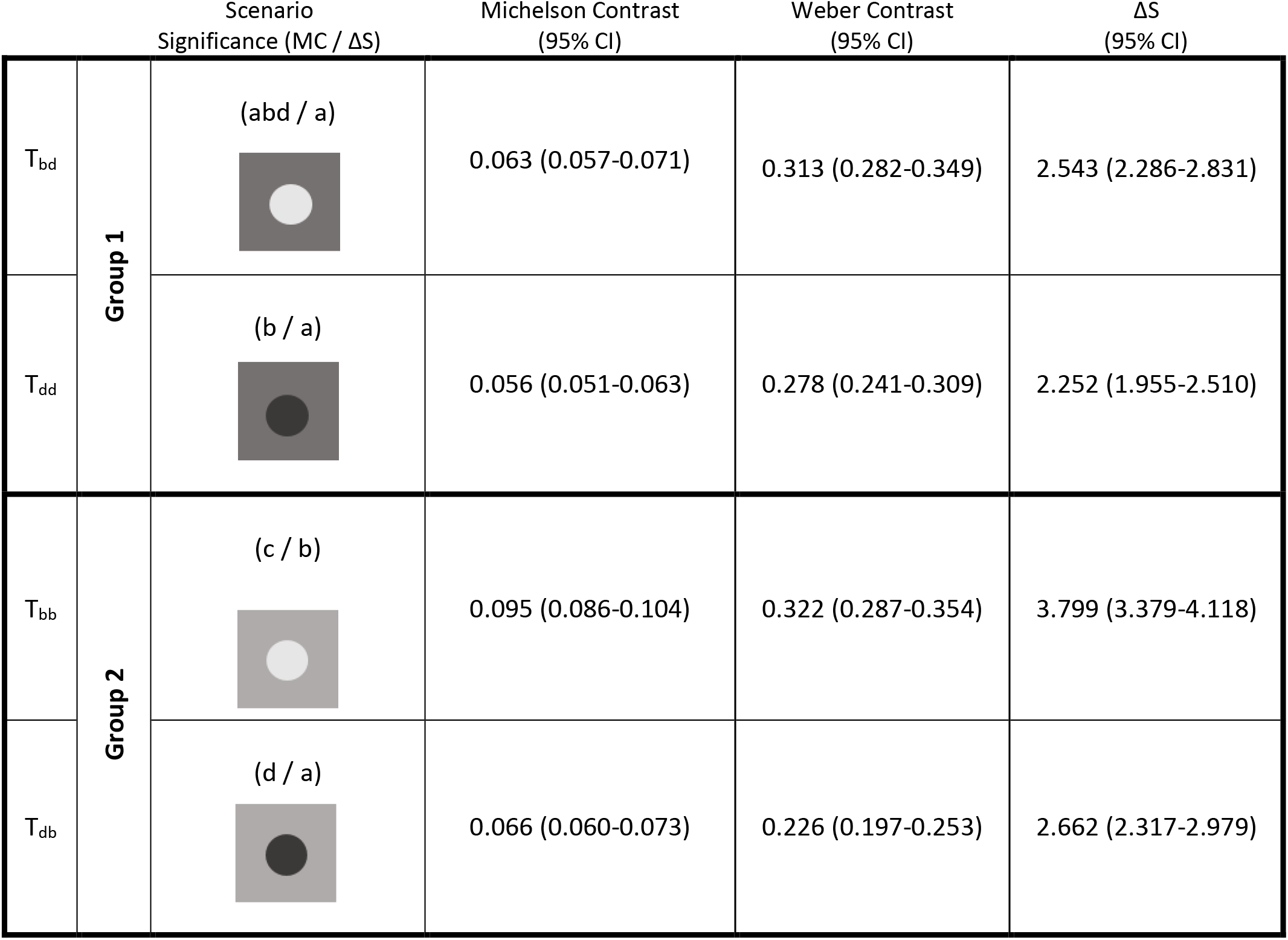
Summary of results for the 50% correct choice threshold contrasts. Letters above scenario drawings indicate significant differences in MC thresholds as per bootstrap sampling (letters on the left) or a 1 ΔS RNL contrast (letters on the right).

|  |  | Scenario<br>Significance (MC / ΔS) | Michelson Contrast<br>(95% CI) | Weber Contrast<br>(95% CI) | ΔS<br>(95% CI) |
| --- | --- | --- | --- | --- | --- |
| T<sub>bd</sub> | Group 1 | (abd / a) | 0.063 (0.057-0.071) | 0.313 (0.282-0.349) | 2.543 (2.286-2.831) |
| T<sub>dd</sub> |  | (b / a) | 0.056 (0.051-0.063) | 0.278 (0.241-0.309) | 2.252 (1.955-2.510) |
| T<sub>bb</sub> | Group 2 | (c / b) | 0.095 (0.086-0.104) | 0.322 (0.287-0.354) | 3.799 (3.379-4.118) |
| T<sub>db</sub> |  | (d / a) | 0.066 (0.060-0.073) | 0.226 (0.197-0.253) | 2.662 (2.317-2.979) |

While the threshold for detecting a bright spot against a dark background was not different from that for detecting a dark spot against a bright background (its ‘inverse’ scenario) (T_bd_ -T_db_ =−0.003, CI_diff_ [−0.013 / −0.016]) all other detection thresholds varied significantly from each other when compared across group 1 & 2 (Fig. 3 & 4, Table 2).

**Figure 4:**
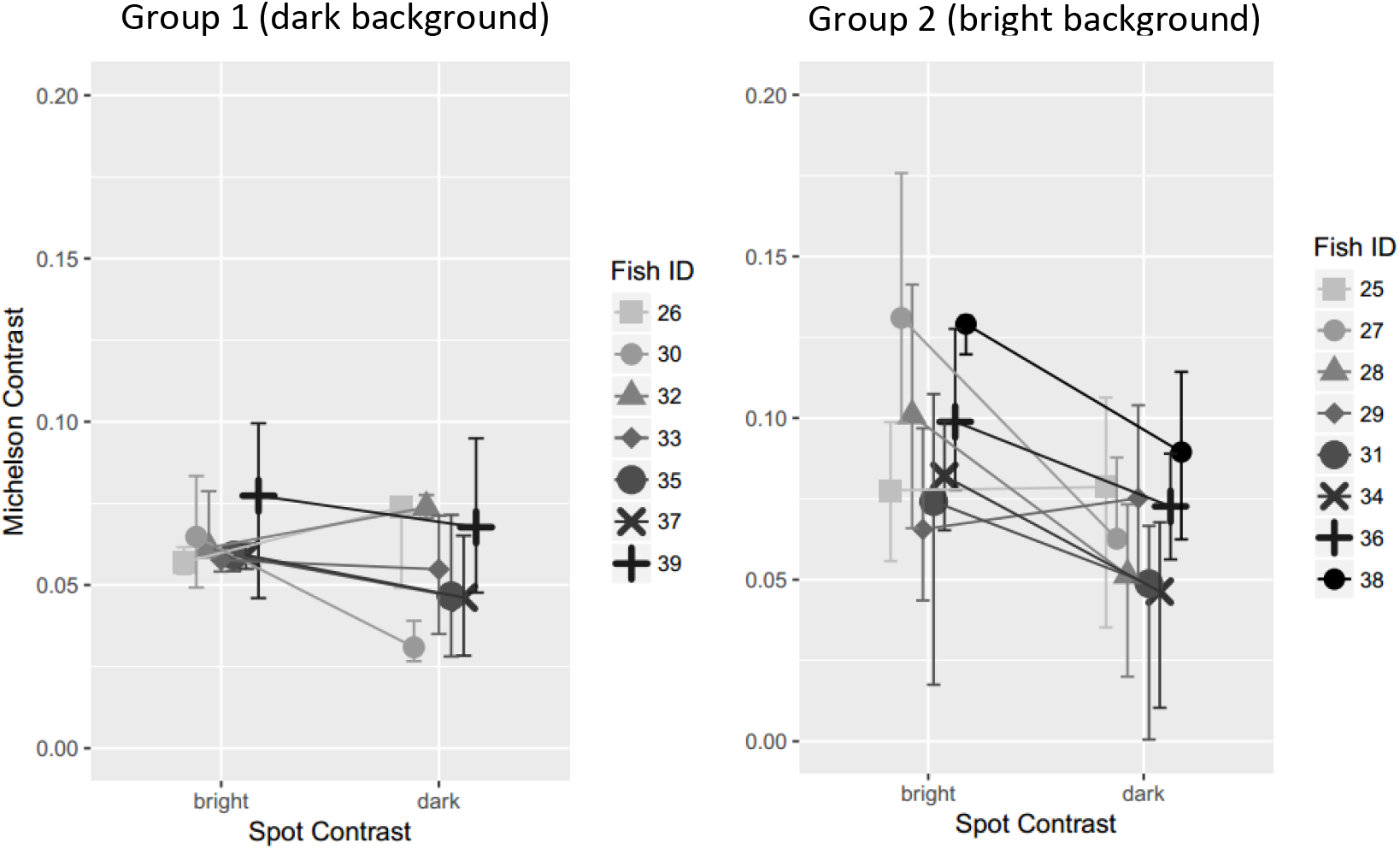
Detection thresholds for individual fish. Individually estimated discrimination thresholds in Michelson contrast for each scenario. Error bars represent the 95% confidence intervals.

## Discussion

Our study demonstrates that for triggerfish, *Rhinecanthus aculeatus*, the ability to discriminate a large, well-illuminated achromatic stimulus against a uniform achromatic background depends on both the relative luminance contrast between target and background (*f*_*t*_ vs. *f*_*b*_) as well as the absolute luminance level (*f*_*t*_ + *f*_*b*_) at which the contrast is perceived (eq. 2). For example, discrimination thresholds, measured as Michelson contrast (MC), were significantly lower when fish were presented with a bright spot against a dark background, as opposed to a bright spot against a bright background (Table 2). However, when expressed in terms of Weber contrast (i.e. scaling the contrast with the luminance level at which the luminance contrast is perceived) these two thresholds were almost identical (Table 2). This finding supports the Weber-Fechner law that states the ability to discriminate a target stimulus against its background scales with the intensity at which the discrimination is made. The same holds true for the discrimination thresholds of dark spot against a dark background (T_dd_) as opposed to a bright background (T_db_), which have an almost identical Weber contrast (Table 2). Furthermore, the contrast sensitivity depends on the direction of the contrast (*f*_*t*_ > *f*_*b*_ ≠ *f*_*t*_ < *f*_*b*_), that is, the Weber contrast for detecting stimuli darker than their respective backgrounds is lower (= more sensitive) from that for stimuli which are brighter than their backgrounds (WC 0.23 – 0.28 for dark spots and 0.31 – 0.32 for bright ones) (Table 2).

Our results agree with previous findings that humans (e.g. Bowen, Pokorny, & Smith, 1989; Emran et al., 2007; Lu & Sperling, 2012), non-human vertebrates (e.g. Baylor et al., 1974), and invertebrate visual systems (e.g. Smithers et al., 2019) are consistently better at detecting darker stimuli. Increasing and decreasing luminance changes are thought to be processed differently: darker stimuli are detected by off-centre ganglion cells, while lighter ones are detected by on-centre ganglion cells (Schiller et al., 1986). Dark stimuli cause depolarization of photoreceptors, whereas light ones are detected as hyperpolarization (Baylor et al., 1974). For example, investigation of turtle photoreceptors has shown that dark stimuli result in much greater depolarization of photoreceptors, than the magnitude of hyperpolarization resulting from light ones (Baylor et al., 1974).This asymmetry is thought to be a crucial contributor to object and motion detection in post-retinal processing (e.g. Oluk et al., 2016; Vidyasagar and Eysel, 2015).

### Behavioural calibration of the RNL

The relationship of absolute (background + stimulus) and relative luminance (background vs. stimulus) contrast does not hold when expressing thresholds as achromatic ΔS (Table 2). The exclusion of signal intensity is a fundamental assumption when calculating chromatic contrasts using the RNL model (Vorobyev and Osorio, 1998), which was designed to quantify contrast perception between two closely opposed chromatic stimuli viewed against an achromatic background. As a result, the RNL equations used by Siddiqi et al. (2004) calculate a relative comparison of two background adapted receptor responses without scaling the difference in photoreceptor stimulation between stimulus and background in relation to the overall brightness of a scene. Thus, the commonly used RNL equations in Siddiqi *et al.* (2004) fail to reflect the Weber-Fechner law for the discrimination of a stimulus from its background. Olsson et al. (2018) proposed the use of an adaptation where the Weber contrast at the behaviourally determined discrimination threshold (*WC*_*t*_) should be used in place of the receptor noise:

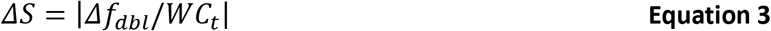

This renders the following ΔS values at threshold: T_dd_ = T_bd_ = 0.41 ΔS ± 0.0001 and T_bb_ = T_db_ = 0.59 ΔS ± 0.001 using the WC determined in this experiment. This makes the RNL model, as modified by Olsson et al. 2018, conform with the Weber-Fechner law while preserving the difference in contrast sensitivity regarding increments and decrements. Furthermore, the thresholds are well below 1 ΔS, making the assumption of a ‘Just Noticeable Difference’ (JND) corresponding to a threshold of 1 ΔS a comfortably conservative (but not extreme) threshold. It should be noted that the general conclusions of Siddiqi et al., (2004) remain most likely correct, but we can now realise a closer description of the underlying mechanisms.

Olsson et al. (2018) propose the use of Michelson contrast (MC) in place of receptor noise in order to estimate the channel specific noise (*e*_*i*_). First, the contrast sensitivity (*CS*) is calculated as the inverse of the behaviourally determined Michelson Contrast (*C*_*t*_):

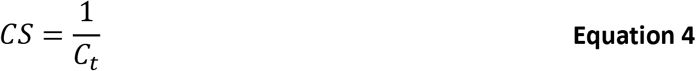

Next, this CS (which is sensitive to the absolute level of luminance as our results confirm) can be used to calculate the relative quantum catch of stimulus 2 (*q*_*stim2*_):

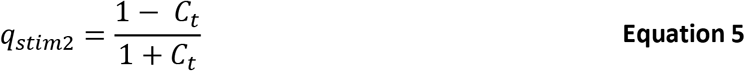

We can then use *q*_*stim2*_ as *f*_*t*_ and our originally measured *f*_*b*_ in eq. 2 to derive the channel noise (*e*_*i*_) (see Olsson et al. 2018 for further details). With the assumption of ΔS = 1 at threshold, this produces:

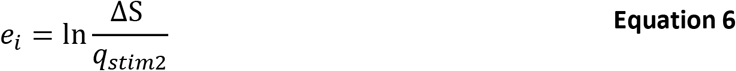

Thus, we obtain the following channel noise estimates (*e*_*i*_ ± 95% CI): *e*_*Tdd*_ = 0.113 (0.098 – 0.125) & *e*_*Tbd*_ = 0.127 (0.114 – 0.142) for group 1 (dark background) and *e*_*Tdb*_ = 0.133 (0.116 – 0.149) & *e*_*Tbb*_ = 0.190 (0.169 – 0.209) for group 2 (bright background). This is the same as setting ω (which is equal to *e*_*i*_) in eq. 1 so that the observed ΔS in our experiments (Table 2) would be equal to 1 (as we determined *f*_*t*_ at threshold by fitting a model to the observed MC). This is interesting, as these noise values are up to almost 4 times as high as the ‘conservative’ (!) standard deviation of noise estimate of 0.05, currently used for modelling vertebrate vision across the field of visual ecology.

Given that WC is meant to be used for comparably small stimuli against large backgrounds and MC to be used for contrasts between stimuli of comparable size, we recommend a differentiated use of either equation 3 or 4–6 depending on the visual context in which a discrimination threshold is used. For example, as the scenario in this study involved the discrimination of a single spot against a much larger background, we would assume equation 3 to be more relevant than equations 4–6 (e.g. equations 4–6 still produce a higher noise ratio for Group 2 (bright background), especially T_bb_). Thus, equations 4-6 would likely be more relevant when discriminating between two objects of equal size. This further implies that one could plot the discrimination curves as a function of WC rather than MC to obtain the discrimination threshold. The thresholds would then only be distinguishable based on the relative direction of the contrast (bright spot or dark spot) and not the background intensity (Table 2). It also implies that thresholds obtained from experiments using a discrimination scenario more fitting to equations 4-6 (e.g. Lind et al., 2013) should not be used to infer the detectability of most likely relatively small prey items against their most likely large visual backgrounds.

### Future directions

The specific mechanisms causing the observed difference in WC between the detection of a dark spot and a bright spot (or mathematical approximations thereof), or an explanation as to why *e*_*achromatic*_ is much higher than the conservatively chosen receptor noise of 0.05, remain speculative. Further investigations might seek advances in the understanding of neurophysiological mechanisms underlying luminance contrast perception in *R. aculeatus*. These include knowledge of the detailed anatomy and receptor noise of double cone photoreceptors, the relative contribution of each double-cone member to luminance contrast sensitivity (Siebeck et al., 2014) as well as the precise mechanism by which photoreceptor stimulation is integrated in post-receptor structures such as edge detecting receptive fields. Behavioural experiments with closely related species with different retinal morphologies would be of interest to further investigate e.g. the role of retinal neuroanatomy on luminance contrast perception.

The adaptations to the RNL model in Olsson et al. (2018), while apparently effective, do not account for the effects of spatial frequency on luminance contrast sensitivity when discriminating objects against visual backgrounds. This is probably the most notable confounding effect on low-level processing of luminance contrast as a result of post-receptor lateral-inhibition (Veale et al., 2017). One possible approach would be the use of contrast sensitivity functions (CSF) to scale Weber fractions as a function of spatial frequency in a visual scene. However, given that these are determined using a perceptually different experimental setup (da Silva Souza et al., 2011) this should be investigated using context specific behavioural experimentation.

Our results warrant caution in the use of uniform contrast sensitivity thresholds (be it achromatic or chromatic) across widely diverse perceptual contexts, independently of which models are used to describe them. Luminance discrimination, as expected, is not just limited by photoreceptor noise and therefore cannot be adequately represented by the use of a singular detection or discrimination threshold determined using the equations in Siddiqi et al. (2004) as currently common in behavioural ecology studies. This realisation shares many parallels with ongoing discussions regarding the use of the RNL model outside of model assumptions (Marshall, 2018; Olsson et al., 2018; Osorio et al., 2017; Sibeaux et al., 2019; Stuart-Fox, 2018; Vasas et al., 2018). Our results suggest the use of conservative achromatic RNL threshold assumption of 3ΔS (e.g. Spottiswoode & Stevens, 2010) without adaptations such as those proposed by Olsson et al. (2018) might warrant caution.

We show that the noise in the achromatic channel of *R. aculeatus* can be substantially higher than anticipated in previous studies modelling its luminance contrast sensitivity using ‘conservative’ receptor noise estimates. However, this increase in channel noise (*e*_*i*_) can be originating from many potential sources, including electrophysiological coupling of receptors in the double cone of *R.aculeatus* (but also a generally higher noise level in receptor responsible for luminance contrast detection) or downstream (post-receptor) processing of visual information. As such it is wrong to conclude receptor noise from such behavioural calibration (Vasas et al., 2018) and it would be more appropriate to refer to the noise of the entire pathway involved in the performance of a task based on the animal’s ability to perceive luminance contrast in a specific visual context.

Despite having investigated luminance contrast sensitivity using two different levels of background luminance, our study only considered discrimination of large, uniform and achromatic circular target stimulus against a uniform grey background. In future studies, more realistic backgrounds and illumination should be taken into account (e.g. Matchette et al., 2020), as a variety of factors can fundamentally influence luminance contrast perception in most circumstances (Gilchrist, 2014; Gilchrist and Radonjic, 2009; Gilchrist et al., 1999; Kingdom, 2011; Maniatis, 2014). Unsurprisingly then, there is evidence that luminance contrast modulates the salience of objects at stages well beyond the retina (Einhäuser and König, 2003).

### Summary

Our findings provide insight into the processing of achromatic information as well as the use of the RNL model to quantify achromatic discrimination by non-human observers. We show that the current use of the RNL model for the quantification of luminance contrast sensitivity thresholds warrants caution. More specifically, our study suggests the lack of adequate scaling of thresholds by the RNL model to the average luminance of a scene and the need for context specific behavioural experimentation whenever possible.

One of the main reasons why researchers use the RNL model is that, presumably, the discriminability of visual contrasts can be reliably predicted by using a set of conservatively estimated physiological parameters such as photoreceptor noise, abundance and spectral sensitivity. While this seems to work satisfyingly well for colour contrast perception across a range of animals, our study suggests quite the opposite to be the case for achromatic contrast. Despite the possibility of calibrating the RNL using contextualised behavioural experiments (as suggested by Olsson et al. 2018), the result remains unsatisfying. However, we recommend the use of behaviourally determined discrimination thresholds suitable to the given visual context in which they are to be applied as well as generous caution when predicting the discriminability of luminance contrast.

Our study indeed suggests that one cannot reliably use the RNL to predict achromatic contrast perception without context specific behavioural experimentation. This has direct implications on the design of behavioural experiments where validated discrimination thresholds are unavailable. For example, given the difficulty of predicting luminance discriminability, luminance contrast should be thoroughly randomised (as opposed to attempting iso-luminance between stimuli) in any behavioural experiment than can potentially be influenced by luminance contrast perception.

## Supporting information

Data and contrasts

Light measurements in tanks

## Contributions

Cedric P. van den Berg conceived and conducted the study and wrote the manuscript. Laurie J. Mitchell, Michelle Hollenkamp and Erin Watson assisted with training and testing of the animals. Karen L. Cheney, Naomi F. Green and N. Justin Marshall helped review the document.

## Acknowledgements

We would like to thank Natalie Meiklejohn for assisting with experiments and data entry, Ama Wakwella for preliminary experimentation and many volunteers for assisting with animal husbandry, Almut Kelber, Anna Hughes & Simon Laughlin for valuable discussions and William Allen & Nick Scott-Samuel for valuable feedback on the manuscript.

